# Competitive advantage of oral streptococci for colonization of the middle ear mucosa

**DOI:** 10.1101/2021.07.28.454259

**Authors:** Kristin M. Jacob, Gemma Reguera

## Abstract

The intermittent aeration of the middle ear seeds its mucosa with saliva aerosols and selects for a distinct community of commensals adapted to the otic microenvironment. We gained insights into the selective forces that enrich for specific groups of oral migrants in the middle ear mucosa by investigating the phylogeny and physiology of 19 strains enriched (*Streptococcus*) or transiently present (*Staphylococcus, Neisseria* and actinobacterial *Micrococcus* and *Corynebacterium*) in otic secretions. Phylogenetic analyses of full length 16S rRNA sequences resolved close relationships between the streptococcal strains and oral commensals as well as between the transient migrants and known nasal and oral species. Physiological functions that facilitate mucosal colonization (swarming motility, surfactant production) and nutrition (mucin and protein degradation) were widespread in all the otic cultivars, as was the ability of most of the isolates to grow both aerobically and anaerobically. However, streptococci stood out for their enhanced biofilm-forming abilities under oxic and anoxic conditions and for their efficient fermentation of mucosal substrates into lactate, a key metabolic intermediate in the otic trophic webs. Additionally, the otic streptococci inhibited the growth of common otopathogens, an antagonistic interaction that could exclude competitors and protect the middle ear mucosa from infections by transient pathobionts. These adaptive traits allow streptococcal migrants to colonize the otic mucosa and grow microcolonies with syntrophic anaerobic partners, establishing trophic webs with other commensals similar to those formed by the oral ancestors in buccal biofilms.

**Importance:** The identification of a diverse microbiome in otic secretions from healthy young adults challenged the entrenched dogma of middle ear sterility and underscored previously unknown roles for oral commensals in the seeding of otic biofilms. By comparing the physiology of novel lineages of streptococci and transient (peri)oral species isolated from otic secretions, we identified adaptive behaviors that allow specific oral streptococcal species to successfully colonize the mucosa of the middle ear. We also describe antagonistic properties of the otic streptococci that help them outcompete transient nasal and oral migrants, including known otopathogens. This knowledge is important to predictively understand the functionality of the otic communities, their interactions with the host mucosa and the outcome of infections.

## Introduction

The oral cavity provides a heterogenous landscape of surfaces and microenvironments (teeth, gingiva, tongue, cheek, hard and soft palate, etc.) for the growth of microbial communities (1). The availability of dietary substrates supports the growth and diversification of the oral inhabitants, making these communities some of the richest and most diverse in the human body (2). Many of these microbes readily disperse with saliva into perioral regions (1) and, from there, to other parts of the digestive tract (3). Dispersal is also enhanced by the spread of saliva aerosols from the oropharynx (back of the throat) into the respiratory tract (3). Indeed, the air inhaled through the nasal passages or the mouth disperses the saliva aerosols from the oropharynx into the esophagus and trachea and, from there, to the lower parts of the aerodigestive tract. Similarly, air exhaled from the lungs disperses saliva aerosols from the oropharynx into the nasal cavity. The opening of the extended tube of the middle ear (the Eustachian tube, Fig. 1) into the distal part of the nasopharynx also promotes the entry of saliva aerosols with exhaled air (4). The Eustachian tube is passively collapsed at rest to sound proof the tympanic cavity and minimize microbial entry, yet it opens when we swallow to draw in air from the lower airways and ventilate the tympanic cavity (Fig. 1) (4). The periodic aperture and collapse of the Eustachian tube promotes the intermittent aeration of the tympanic cavity, relieves negative pressure across the eardrum and secretes into the nasopharynx excess mucus and fluids (4). It also introduces saliva aerosols into the middle ear and establishes fluctuating redox conditions optimal for the growth of strict and facultative anaerobes (5). Consistent with this, otic secretions collected at the orifice of the Eustachian tube are enriched in obligate or facultatively anaerobic genera within the phyla Bacteroidetes (*Prevotella* and *Alloprevotella*), Fusobacteria *(Fusobacterium* and *Leptotrichia*) and Firmicutes (*Veillonella* and *Streptococcus*) (5). The co-enrichment in otic secretions of Bacteroidetes, streptococci and *Veillonella* spp. suggests that they grow syntrophically as in oral biofilms (5). In this model (Fig. 1), Bacteroidetes could degrade mucin glycoproteins and other mucosal proteins into substrates (sugars and peptides) that oral streptococci can ferment under both aerobic and anaerobic conditions (6). This metabolic capacity could produce lactate for *Veillonella* fermentation into propionate, acetate, CO_2_ and H_2_, as described in oral biofilms (7, 8). The lactate dependency of *Veillonella* spp. could also support metabolic interactions with Bacteroidetes partners that ferment simple sugars into lactate (9). The co-aggregative properties of oral streptococci during the formation of the dental plaque (2, 10) could facilitate the formation of microcolonies in the middle ear as well and promote metabolic exchange with the strictly anaerobic Bacteroidetes and *Veillonella* partners (5). Through their collective metabolism (Fig. 1), otic microcolonies of Bacteroidetes, streptococci and *Veillonella* could degrade and ferment host-derived mucins and proteins into short chain fatty acids such as propionate and acetate, potentially contributing to mucosal health as in other body sites (11).

**Fig 1:**
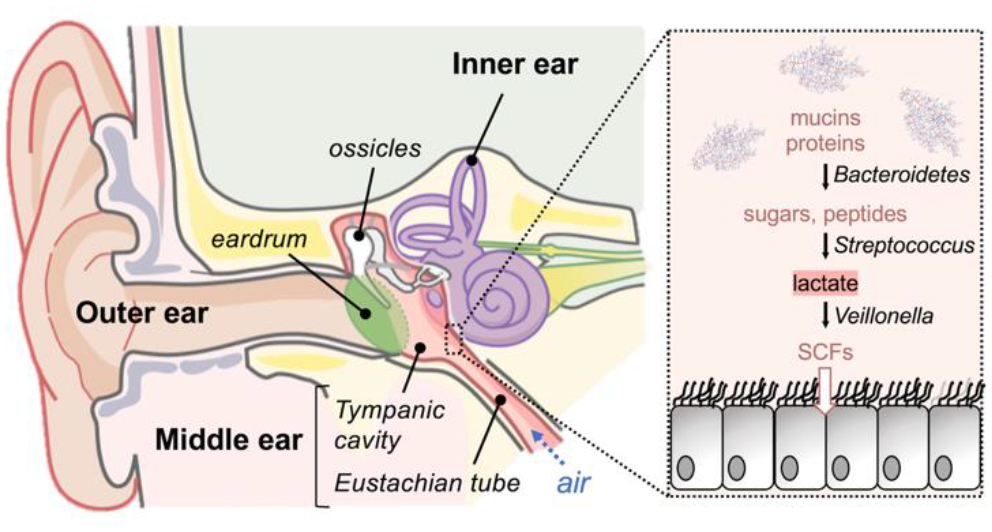
Illustration of the human ear anatomy (*left*) and trophic webs within bacterial microcolonies in the middle ear mucosa (*right*). The human ear is divided in three compartments (outer, middle and inner). The eardrum separates the outer ear canal from the tympanic cavity of the middle ear, which extends as a tube (Eustachian tube) into the nasopharynx to draw in air and drain otic secretions. The microbiome sequenced from otic secretions of healthy young adults (5) supports the establishment of an otic trophic web (inset) for the degradation of host mucins and proteins by Bacteroidetes into substrates (sugars and peptides) that *Streptococcus* and *Veillonella* cooperatively ferment into short chain fatty acids (SCFs) via lactate.

The presence of bacterial microcolonies in biopsy specimens of the mucosal lining of the tympanic cavity (12, 13) suggests that some oral migrants penetrated the mucus layer, attached to the underlying epithelium and formed biofilms. Swarming motility could have allowed these successful colonizers to move through the viscous mucus medium and evade immune attack (14). Crossing the mucus barrier is also important to avoid clearance, which in the middle ear is mediated by the movement of cilia (mucociliary clearance) and the pumping force exerted by muscles contracting and relaxing around the Eustachian tube when swallowing (muscular clearance) (15, 16). Once they reach the mucosal epithelium, successful colonizers attach to it and grow as microcolonies on the mucosal surface to avoid clearance (17). This points at biofilm formation as a critical selective factor for growth and reproduction in the otic mucosa. Biofilms are also important to create anoxic microenvironments for anaerobic partners to grow despite the periodic redox fluctuations experienced in the middle ear. Because the aperture of the Eustachian tube is triggered by swallowing, air only enters the middle ear every minute during the wake hours or every five minutes during sleep (4). Facultative anaerobes such as *Streptococcus* have a growth advantage under these conditions, and could co-aggregate as in oral biofilms (18) to establish anaerobic niches for their anaerobic syntrophic partners (e.g., Bacteroidetes and *Veillonella* species). Nutrient foraging is also important for the establishment of biofilms in the middle ear mucosa. In the absence of dietary nutrients, the otic communities are more likely to sustain their trophic webs with host-derived nutrients such as proteins and mucin glycoproteins secreted to the mucosa (19). Motility in the viscous environment of the mucosa could facilitate nutrient foraging (20) while the ability of the bacteria to secrete hydrolytic enzymes (mucinases and proteases) could allow them to break down the mucin glycoproteins and mucosal proteins into sugars and amino acids (21).

Phylogenetic analyses of cultivars recovered from otic secretions also support an oral ancestry of the middle ear communities (5). These cultivars include streptococcal strains closely related yet phylogentically distinct from oral ancestors, consistent with the diversification of oral taxa into lineages better suited for growth and reproduction in the middle ear mucosa (5). Cultivars recovered from otic secretions also include nasal and oral species from genera (*Staphylococcus, Neisseria* and actinobacterial *Micrococcus* and *Corynebacterium*) that transiently disperse through the oral and perioral regions (5). Streptococcal and staphylococcal species are, for example, among the most prominent members in the oral and nasal microbiomes, respectively (3, 22). Both groups disperse in the aerodigestive tract and are predicted to enter the middle ear during the intermittent cycles of aperture of the Eustachian tube. Yet, while streptococci are one of the most abundant groups in otic secretions, staphylococcal-like sequences are seldom detected (5). This suggests that streptococcal migrants have a competitive advantage over the transient staphylococcal species during the colonization of the middle ear mucosa. To test this, we sequenced and partially assembled the genomes of 19 cultivars representing otic commensals (*Streptococcus*) and transient genera (*Staphylococcus, Neisseria, Micrococcus* and *Corynebacterium*) (5) and used the full length 16S rRNA sequences to identify their closest relatives. We then screened the cultivars for adaptive traits predicted to be important for mucosal colonization (swarming, surfactant production, biofilm formation) and for growth under conditions (redox, nutritional, etc.) relevant to the middle ear microenvironment. Our study revealed similar adaptive traits for mucosal growth in most of the isolates but aggregative and metabolic properties of streptococci that could facilitate their attachment to the mucosal epithelium and syntrophic growth within microcolonies. These same properties are also present in their closest oral ancestors, with whom they share the ability to establish trophic webs with anaerobes and antagonize transient members from the oral and perioral regions. These findings provide novel insights into the adaptive responses that sustain the growth and functionality of otic communities as well as interactions with the host mucosa and transient migrants that influence the outcome of infections.

## Results

### Phylogenetic analysis supports the oral ancestry of otic streptococcal commensals

Cultivars recovered from otic secretions collected from healthy young adults include genera enriched in the otic secretions (*Streptococcus*) as well as transient or lowly abundant groups (*Staphylococcus, Neisseria,* and the actinobacterial genera *Micrococcus* and *Corynebacterium*) (5). Phylogenetic analysis of partial 16S rDNA sequences amplified from these isolates revealed close relationships with oral (oropharyngeal and buccal) strains isolated from the same individuals but lacked the resolution needed for species-level demarcation (5). Thus, we sequenced and partially assembled the genomes of 19 otic cultivars to retrieve full-length 16S rDNA sequences for each of the isolates. A species sequence identity cutoff of >98.7% (23) matched each otic isolate to more than one species within each genus (Table 1 shows the top identity hit for each strain). Phylogenetic inference methods resolved, however, close evolutionary ties with specific species that reside or are transiently isolated from the oral cavity (Fig. 2).

**Fig. 2:**
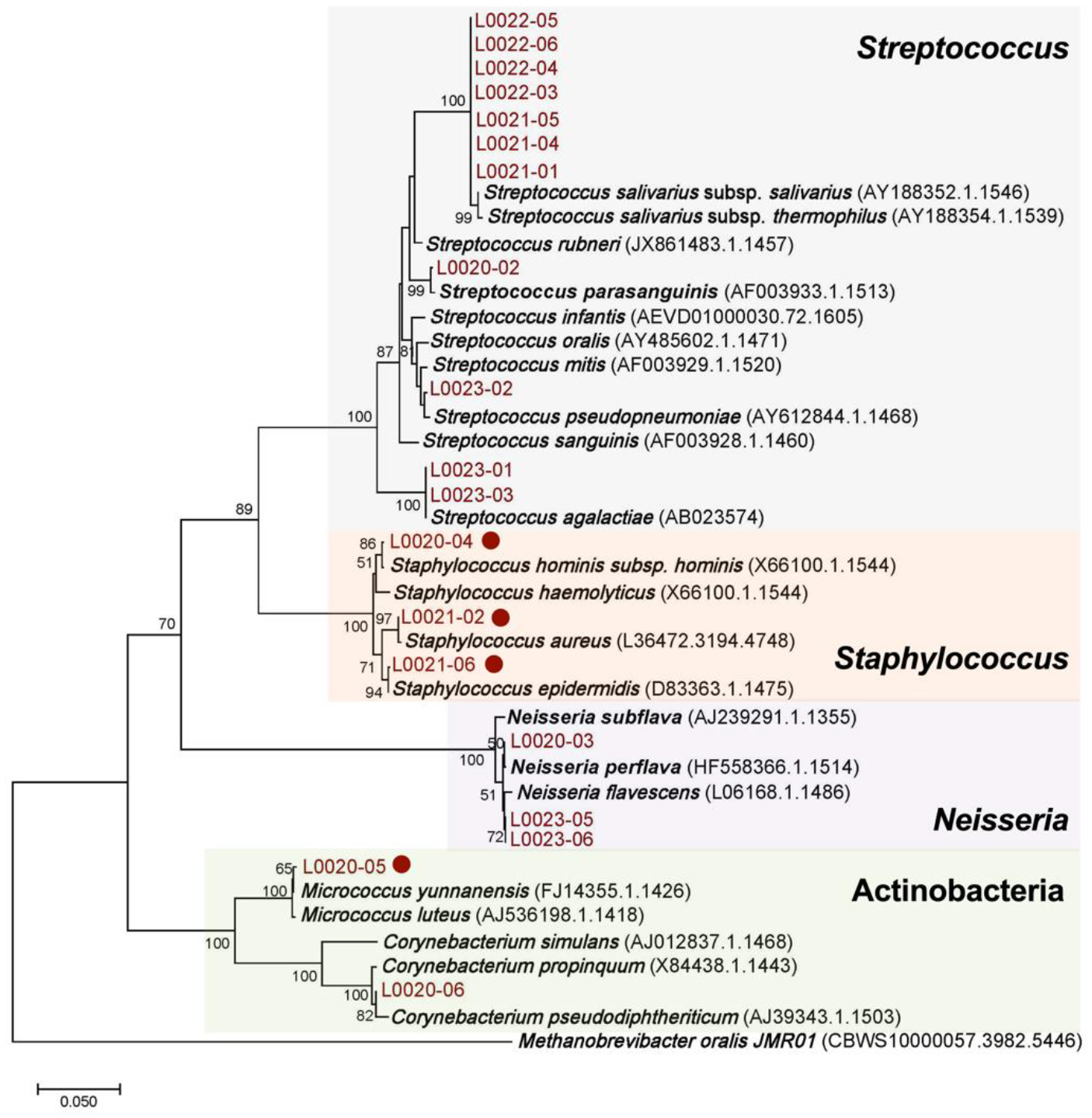
16S rRNA gene phylogeny of otic cultivars. Maximum-likelihood tree constructed with full-length 16S rRNA sequences from the otic isolates and the closest reference strains (accession number in parentheses). The scale bar indicates 5% sequence divergence filtered to a conservation threshold above 79% using the Living Tree Database (24, 25). Bootstrap probabilities by 1000 replicates at or above 50% are denoted by numbers at each node. The circles identify catalase-positive isolates.

**Table 1:** Taxonomic classification (reference strain) of otic strains based on the % identity (ID) of their full-length 16S rRNA sequence.

| Strain | GenBank no. | Reference Strain (Accession; % ID) |
| --- | --- | --- |
| *Streptococcus* |  |  |
| L0020-02 | MW866489 | *Streptococcus parasanguinis* (NR_024842.1; 99.47) |
| L0021-01 | MW866494 | *Streptococcus salivarius* (NR_042776.1; 99.81) |
| L0021-04 | MW866496 | *Streptococcus salivarius* (NR_042776.1; 99.81) |
| L0021-05 | MW866497 | *Streptococcus salivarius* (NR_042776.1; 99.81) |
| L0022-03 | MW866499 | *Streptococcus salivarius* (NR_042776.1; 99.81) |
| L0022-04 | MW866500 | *Streptococcus salivarius* (NR_042776.1; 99.81) |
| L0022-05 | MW866501 | *Streptococcus salivarius* (NR_042776.1; 99.81) |
| L0022-06 | MW866502 | *Streptococcus salivarius* (NR_042776.1; 99.81) |
| L0023-01 | MW866503 | *Streptococcus agalactiae* (NR_040821.1; 100) |
| L0023-02 | MW866504 | *Streptococcus oralis* (NR_117719.1; 99.47) |
| L0023-03 | MW866505 | *Streptococcus agalactiae* (NR_040821.1; 100) |
| *Staphylococcus* |  |  |
| L0020-04 | MW866491 | *Staphylococcus hominis* (NR_036956.1; 99.61) |
| L0021-02 | MW866495 | *Staphylococcus aureus* (NR_037007.2; 99.87) |
| L0021-06 | MW866498 | *Staphylococcus saccharolyticus* (NR_113405.1; 99.4) |
| *Micrococcus* |  |  |
| L0020-05 | MW866492 | *Micrococcus luteus* (NR_075062.2, 99.61) |
| *Corynebacterium* |  |  |
| L0020-06 | MW866493 | *Corynebacterium pseudodiphthericum* (NR_042137.1; 99.47) |
| *Neisseria* |  |  |
| L0020-03 | MW866490 | *Neisseria perflava* (NR_114694; 99.93) |
| L0023-05 | MW866507 | *Neisseria perflava* NR_117694.1; 99.74) |
| L0023-06 | MW866506 | *Neisseria perflava* NR_117694.1; 99.74) |

The nearest neighbor to most of the *Streptococcus* sequences (seven of them) was *Streptococcus salivarius* (subspecies *salivarius* and *thermophilus*) (Fig. 2). Genomic divergence (size and gene content) is high for species and subspecies within the Salivarius group (26). As a result, strains of *S. salivarius* can have very different metabolic and physiological characteristics or even habitat/host preferences despite high 16S rRNA sequence identity (26, 27). Thus, the separate clustering of the 7 otic strains of *S. salivarius* could reflect substantial divergence of the otic subclade from an oral ancestor. The rest of the otic streptococcal sequences clustered separately from close oral relatives within the Mitis (L0020-02 and *Streptococcus parasanguinis*), Viridans (L0023-02 and *Streptococcus pseudoneumoniae*) and the Lancefield’s group B streptococcus or GBS (L0023-01 and L0023-03 and *Streptococcus agalactiae*) groups (27, 28) (Fig. 2). Hence, 16S rRNA phylogeny supports the oral ancestry of the streptococcal cultivars but also reveals a level of divergence that is consistent with the ecological diversification of niche-adapted otic lineages proposed in earlier studies (5).

The 16S rRNA sequence identity of the non-streptococcal strains also produced more than one match to species of *Staphylococcus, Neisseria, Micrococcus* and *Corynebacterium* (Table 1). Consistent with the classification of three of the isolates as *Staphylococcus* spp., all were catalase negative, and all branched within subclades of staphylococcal 16S rRNA sequences (Fig. 2). The closest neighbors to the three otic staphylococci were species (*Staphylococcus hominis, Staphylococcus aureus*, and *Staphylococcus epidermidis*) that are highly represented in the nasal passages (22). Their nasal abundance facilitates their dispersal in the contiguous oral cavity (29) and their transient detection in oral and perioral regions (29, 30), including otic secretions (5). Transience also explains the recovery of *Neisseria* strains closely related to *Neisseria perflava, Neisseria subflava* and *Neisseria flavescens* (Fig. 2), which are species that colonize the mucosa of the oropharynx (31, 32) and disperse via saliva aerosols (33). Neutral community models predict, however, a similar distribution of *Neisseria* species in otic secretions compared to oral and perioral source communities, supporting the idea that they are transient migrants (5). The otic isolates also included two actinobacterial *Micrococcus* and *Corynebacterium* strains (Table 1). The *Micrococcus* isolate was catalase-positive, a general phenotypic trait of the genus (34), and branched closely to *Micrococcus yunnanensis* (Fig. 2). This is a soil *Micrococcus* species (35) that, like other environmental micrococci, transiently disperses with air in the human aerodigestive tract (36). The second actinobacterial isolate was closely related to *Corynebacterium pseudodiphtericum* (Fig. 2). *Corynebacterium* commensals are prominent members of the nasal microbiomes and antagonists of nasal pathobionts, including some of the most important otopathogens (37). Their abundance in the nasal microflora explains their detection in oral and perioral regions (38). However, actinobacteria only account for ∼1% of the operational taxonomic units (OTUs) in otic secretions, suggesting they are negatively selected for growth and reproduction in the middle ear mucosa (5).

### Surfactant-mediated swarming motility is widespread among the otic cultivars

Successful colonization of any respiratory mucosa requires bacterial migrants to move rapidly across the mucus layer in order to avoid immune attack and clearance (17). Some flagellated bacteria can reach the underlying epithelial lining by rapidly swarming in groups through the viscous mucoid layer, a process that is stimulated by the lubricating effect of mucin glycoproteins (20). Swarming behaviors can be identified in laboratory plate assays that test the expansion of microcolonies on a soft agar (0.4-0.5%) surface (20). Thus, we tested the ability of the 19 otic isolates to swarm on the surface of 0.5% tryptone soy agar (TSA) plates in reference to the robust swarmer *Pseudomonas aeruginosa* PA01 (39). Figure 3A shows the progression of the swarming expansion for each strain over time, which we calculated as the average expansion zone in triplicate plate assays (Table 2). Although *P. aeruginosa* showed large zones of swarming expansion already at 18 h, we only detected swarming activity in the otic isolates after 42 or 62 h of colony growth (Fig. 3A). Lag phases are not unusual prior to swarming on agar plates for cells that must reprogram their physiology to grow on the surface of the agar-solidified medium (20). Consistent with this, the strains that grew faster on the semisolid TSA plates (three staphylococcal and the two actinobacterial isolates) produced visible zones of swarming expansion at 42 h, while the slowest growers (*N. perflava* (L0023-05 and L0023-06) required 62 h of incubation (Table 2). Notably, most of the streptococci grew well in tryptone soy broth (TSB), yet they aggregated strongly when growing on the surface of soft-agar plates (Fig. 3B). These aggregative strains also had delayed swarming phenotypes on 0.5% TSA plates (Table 2) but it was possible to rescue the swarming delay on plates with lower (0.4%) agar concentration (Fig. 3B). For example, the streptococcal strain L0022-03 did not swarm on 0.5% TSA plates until after 62 h (Table 2) but expanded 0.28 cm away from the edge of the macrocolony after 42 h of growth on 0.4% TSA plates (Fig. 3B). Temperate swarmers often require a softer agar surface to overcome frictionally forces between the cell and the surface (20). This is because lowering the agar concentration facilitates water movement to the surface and immerses the cells in a layer of liquid that stimulates swarming (20).

**Fig. 3.**
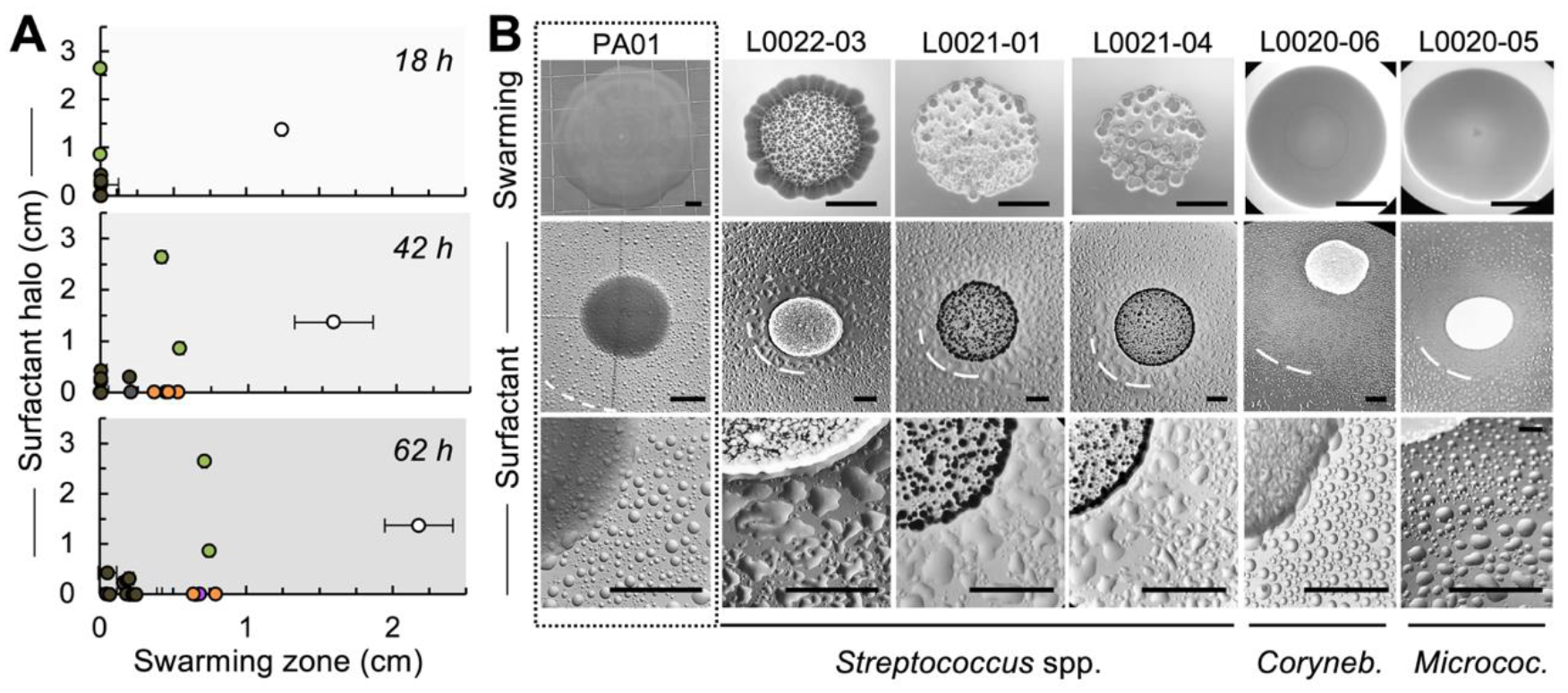
Swarming motility and surfactant production by otic cultivars in reference to *Pseudomonas aeruginosa* PA01. **(A)** Average surfactant production (halo of mineral oil dispersal around 24h colonies grown on 1.5% TSA) and size of swarming expansion (0.5% TSA plates at 18, 42 and 62 h) measured in triplicate replicates of the otic isolates (*Streptococcus* in gray; *Staphylococcus* in orange; *Neisseria* in purple; and actinobacterial strains of *Corynebacterium* and *Micrococcus* in green) and the positive control (*P. aeruginosa* PA01 in white). **(B)** Representative images of swarming (0.4% TSA, 42 h) and surfactant (1.5% TSA, 24 h) plate assays for *P. aeruginosa* PA01 (positive control, boxed) and otic strains of *Streptococcus, Corynebacterium* and *Micrococcus* (scale bars, 0.5 cm). The edge of the surfactant halo is highlighted with a dashed white line.

**Table 2.** **Coaggregation, swarming motility and surfactant production of otic isolates in** reference to positive control (*P. aeruginosa* PA01).

| Species/closest relative | Strain | Aggregation<sup>a</sup> | Surfactant<sup>b</sup> | Swarming<sup>c</sup> |  |  |
| --- | --- | --- | --- | --- | --- | --- |
|  |  |  |  | 18 h | 42 h | 62 h |
| *Streptococcus* |  |  |  |  |  |  |
| *S. parasanguinis* | L0020-02 | + | – | – | – | 0.07 (0.10) |
| *S. salivarius* | L0021-01 | + | 0.23 (0.19) | – | – | 0.16 (0.05) |
| *S. salivarius* | L0021-04 | + | 0.46 (0.09) | – | – | 0.05 (0.12) |
| *S. salivarius* | L0021-05 | + | – | – | – | 0.05 (0.07) |
| *S. salivarius* | L0022-03 | + | 0.28 (0.19) | – | – | 0.19 (0.01) |
| *S. salivarius* | L0022-04 | + | – | – | – | 0.20 (0.08) |
| *S. salivarius* | L0022-05 | + | 0.18 (0.12) | – | – | 0.20 (0.03) |
| *S. salivarius* | L0022-06 | + | 0.09 (0.06) | – | – | 0.18 (0.02) |
| *S. agalactiae* | L0023-01 | + | – | – | 0.21 (0.22) | 0.23 (0.02) |
| *S. pseudopneumoniae* | L0023-02 | + | – | – | – | 0.07 (0.10) |
| *S. agalactiae* | L0023-03 | + | 0.31 (0.11) | – | 0.19 (0.01) | 0.20 (0.06) |
| *Staphylococcus* |  |  |  |  |  |  |
| *S. hominis* | L0020-04 | – | – | – | 0.37 (0.01) | 0.63 (0.08) |
| *S. aureus* | L0021-02 | – | – | – | 0.53 (0.01) | 0.79 (0.01) |
| *S. epidermidis* | L0021-06 | – | – | – | 0.46 (0.03) | 0.64 (0.02) |
| *Actinobacteria* |  |  |  |  |  |  |
| *M. yunnanensis* | L0020-05 | – | 0.86 (0.13) | – | 0.54 (0.05) | 0.75 (0.04) |
| *C. pseudodiphthericum* | L0020-06 | – | 2.64 (0.05) | – | 0.41 (0.06) | 0.72 (0.03) |
| *Neisseria* |  |  |  |  |  |  |
| *N. perflava* | L0020-03 | – | – | – | 0.44 (0.03) | 0.66 (0.01) |
| *N. flavescens* | L0023-05 | + | – | – | – | 0.23 (0.32) |
| *N. flavescens* | L0023-06 | + | – | – | – | 0.25 (0.35) |
| *P. aeruginosa* | PA01 | – | 1.37 (0.06) | 1.24 (0.35) | 1.58 (0.53) | 2.18 (0.65) |

<sup>a</sup> Aggregative (+) or uniform (–) growth of cultures spotted on 0.5% TSA plates.
<sup>b</sup> Average (and standard deviation) of triplicate surfactant haloes (cm) measured as the zone of mineral oil dispersion around colonies grown at 37°C on 1.5% TSA plates. (–, not detected).
<sup>c</sup> Average (and standard deviation) of triplicate swarming expansion zones (cm) around colonies grown at 37°C on soft agar (0.5%) TSA plates for 18, 42 and 62 h. (–, not detected).

The need for some bacteria to express cellular components (flagella, exopolysaccharide, surfactants, etc.) mediating swarming on semisolid agar can also delay the appearance of expansion zones (20). Surfactants are particularly important to reduce frictional resistance between the surface of swarming cells and the underlying substratum (20). Furthermore, their concentration and diffusion in soft-agar medium controls the extent of swarming expansion (40). Hence, we also screened the cultivars for surfactant production. To do this, we spot-plated each isolate on hard agar (1.5%) TSA plates and airbrushed a fine mist of mineral oil droplets onto colonies grown at 37°C for 24 h. This atomized oil assay instantaneously reveals halos of oil droplet dispersal around surfactant-producing strains and allows for semiquantitative estimation of the levels of surfactant production, even when present at concentrations too low to be detected by traditional methods such as the water drop collapse assay (40). We detected haloes of oil dispersal around 9 of the isolates at 24 h (Table 2). Furthermore, we observed positive correlations between surfactant production and swarming ability on 0.5% TSA for most strains (Fig. 3A). For example, the actinobacterial isolates, which were the most robust swarmers, produced the highest levels of surfactant (Table 2). By contrast, temperate swarmers such as the streptococcal isolates produced low or undetectable levels of surfactants under the experimental conditions. As an exception, the staphylococcal isolates swarmed robustly on the soft agar plates (Fig. 3) although they did not produce detectable halos of mineral oil dispersion (Table 2). Staphylococcal cells lack flagellar locomotion and thus do not have canonical (i.e., flagella-driven) swarming behaviors. These bacteria can however passively ‘spread’ on soft agar surfaces (41) through the coordinated synthesis of lubricating peptides known as phenol-soluble modulins (PSMs) (42). PSM surfactants accumulate very close to the colony edge (43). Hence, they are unlikely to produce a halo of oil dispersal in the atomized assay used for testing.

### Redox and nutritional advantage of otic streptococci in the middle ear mucosa

Successful colonizers of the middle ear mucosa face sharp redox fluctuations due to the brief (400 milliseconds) and infrequent (approximately every minute when we swallow) aperture of the Eustachian tube (4). The enrichment in otic secretions of anaerobic metabolisms further supports the idea that conditions of oxygen limitation prevail in the middle ear (5). For this reason, we tested the ability of the otic cultivars to grow under aerobic or anaerobic conditions (Fig. 4A). All the isolates grew well in oxic and anoxic TSB medium, except for two *Neisseria* strains (L0023-05 and L0023-06) that grew slowly in the oxic broth. These two strains also flocculated extensively in oxic medium, an aggregative behavior exhibited by microaerophiles in response to elevated (and toxic) oxygen concentrations (44). By contrast, the streptococcal and staphylococcal strains had similar growth rates aerobically and anaerobically (0.56±0.23 and 0.49±0.12 doubling times, respectively), suggestive of a competitive advantage for growth and reproduction under sharp redox fluctuations. The actinobacterial strains also grew aerobically and anaerobically but differed in their redox preference. Both isolates doubled every ∼0.5 h under anoxic conditions. However, generation times increased in aerobic cultures of *Micrococcus* L0020-05 (∼0.7 h) whereas *Corynebacterium* L0020-06 grew more rapidly than any other strain under these conditions (0.15 h average generation time). This aerobic preference matches well the enrichment of *Corynebacterium* species in the aerated nasal passages (22) and the reduced abundance of this group in otic secretions (5).

**Fig. 4:**
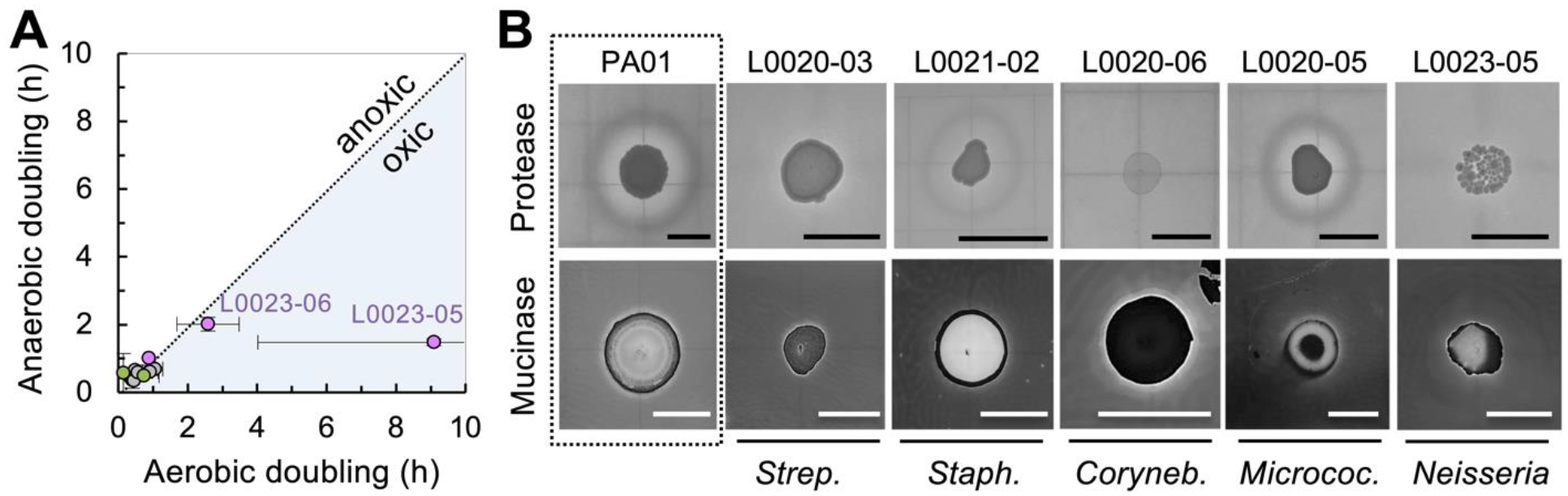
Growth of otic isolates as a function of oxygen availability and host nutrients (protein and mucin). **(A)** Average doubling times of otic isolates growing in triplicate TSB cultures aerobically or anaerobically at 37°C. Data points are color-coded for *Streptococcus* (gray), *Staphylococcus* (orange), *Neisseria* (purple) and actinobacterial genera *Micrococcus* and *Corynebacterium* (green). The flocculating strains of *Neisseria* are labeled. **(B)** Protease and mucinase activity (haloes of milk casein or porcine gastric mucin degradation, respectively) of representative otic isolates and *P. aeruginosa* PA01 (positive control, boxed). The milk casein plates were photographed without staining after 24 h of incubation at 37°C. The mucin plates were incubated for 48 h and stained with 0.1% amido black prior to photography. Scale bars, 0.5 cm.

In addition to redox fluctuations, bacteria colonizing the otic mucosa must cope with a scarcity of nutrients. The limited carriage of dietary substrates in saliva aerosols reduces nutrient availability in the middle ear, exerting selective pressure on colonizers to use host-derived nutrients such as mucosal proteins and mucin glycoproteins (5). To test this, we screened the otic isolates for their ability to secrete enzymes (proteases and mucinases) needed to break down the host nutrients into readily assimilated substrates (amino acids and sugars). For these experiments, we spot-plated the cultivars onto TSA plates supplemented with 5% lactose-free skim milk (protease assay) or 0.5% porcine gastric mucin (mucinase assay) for 24 h to identify zones of substrate degradation around the colonies. Figure 4B shows typical results for representative otic strains and the positive control *P aeruginosa* PA01, while Table 3 shows the results (presence or absence of a halo of substrate degradation) in triplicate plate assays for each strain. Notably, all the isolates were able to degrade mucin to various degrees after 24 h of incubation at 37°C. Three aggregative strains of *S. salivarius* (L0021-01, L0022-03 and L0022-04) produced only faint mucin clearings after 24 h (+/– in Table 3) but the zone of degradation became more prominent after extending the incubation period to 48 h. By contrast, protease activity was only detected in the streptococcal and staphylococcal groups (Table 3). Extracellular proteases typically have low substrate selectivity and, thus, can cleave a wide range of substrates (45). Non-selectivity is particularly advantageous in the middle ear mucosa, as it allows residents to scavenge proteins and the mucin protein backbone as nitrogen sources (19). In addition to providing a metabolic advantage, proteases facilitate mucosal penetration, control mucus viscosity, modulate host immune responses, and can prevent the establishment of competitors (46). Therefore, protease secretion may confer on streptococcal and staphylococcal migrants a competitive advantage for growth and reproduction in the otic mucosa.

**Table 3.**
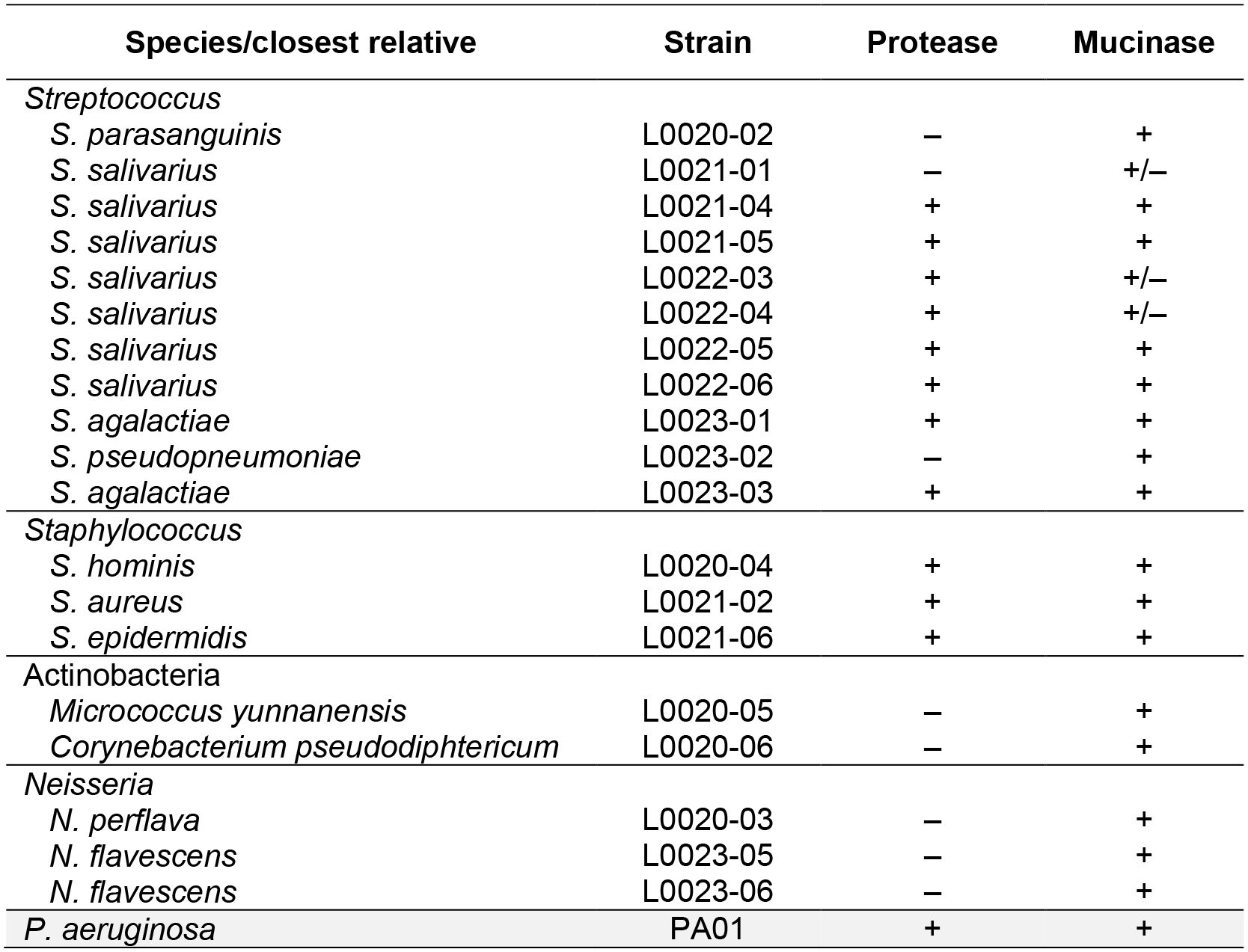
Protease and mucinase enzymatic activity of otic isolates in reference to positive control (P. aeruginosa PA01). Presence (+) or absence (–) of a halo of degradation in TSA plates supplemented with skim milk (protease assay) or mucin (mucinase assay) after 24 h of growth. The presence of a faint halo is indicated with “+/–“.

| Species/closest relative | Strain | Protease | Mucinase |
| --- | --- | --- | --- |
| *Streptococcus* |  |  |  |
| *S. parasanguinis* | L0020-02 | – | + |
| *S. salivarius* | L0021-01 | – | +/- |
| *S. salivarius* | L0021-04 | + | + |
| *S. salivarius* | L0021-05 | + | + |
| *S. salivarius* | L0022-03 | + | +/- |
| *S. salivarius* | L0022-04 | + | +/- |
| *S. salivarius* | L0022-05 | + | + |
| *S. salivarius* | L0022-06 | + | + |
| *S. agalactiae* | L0023-01 | + | + |
| *S. pseudopneumoniae* | L0023-02 | – | + |
| *S. agalactiae* | L0023-03 | + | + |
| *Staphylococcus* |  |  |  |
| *S. hominis* | L0020-04 | + | + |
| *S. aureus* | L0021-02 | + | + |
| *S. epidermidis* | L0021-06 | + | + |
| *Actinobacteria* |  |  |  |
| *Micrococcus yunnanensis* | L0020-05 | – | + |
| *Corynebacterium pseudodiphthericum* | L0020-06 | – | + |
| *Neisseria* |  |  |  |
| *N. perflava* | L0020-03 | – | + |
| *N. flavescens* | L0023-05 | – | + |
| *N. flavescens* | L0023-06 | – | + |
| *P. aeruginosa* | PA01 | + | + |

### Metabolic advantage of streptococci for syntrophic growth in biofilms

The presence of bacterial microcolonies on the epithelial surface of biopsy specimens collected from the tympanic cavity of healthy individuals (13) points at biofilm formation as an important adaptive response for successful colonization of the middle ear mucosa. Based on this, we investigated the ability of the otic isolates to form biofilms under aerobic and anaerobic conditions. These assays used crystal violet to stain 24-h biofilms formed at the bottom of the microtiter plates that we previously used for planktonic growth studies (Fig. 4A). We then solubilized the biofilm-associated dye to estimate the biofilm biomass from the absorbance of the solution at 550 nm (Fig. 5A). All but two streptococcal strains (*S. pseudopneumoniae* L0023- 02 and *S. agalactiae* L0023-03) formed robust biofilms under aerobic conditions. Among these streptococcal biofilm formers, four strains (*S. salivarius* L0021-04 and L0021-05, *S. parasanguinis* L0020-02, and *S. agalactiae* L0023-01) clustered separately with a staphylococcal isolate (*S. aureus* L0021-02) for their ability to also form robust biofilms in anoxic media (Fig. 5A). This contrasts with strains of *S. salivarius* (L0021-01, L0022-03, L0022-04, L0022-05 and L0022-06) that had a biofilm growth advantage in oxic medium only. The enhanced biofilm abilities of these isolates correlated well with pH drops below 5 in the culture broth (Fig. 5B) due to the accumulation of lactate as a fermentation byproduct (*p*=0.03) (Fig. 5C). Culture acidification likely triggered biofilm formation in the streptococcal cultures, because cells entered stationary phase (0.62±0.05 OD_600_) once the pH dropped below 5. These results point at lactate accumulation and broth acidification as the trigger of the planktonic-to-biofilm transition in the otic streptococci. This response is similar to that described for oral streptococcal commensals, which also produce lactic acid as the main fermentation byproduct (47) and stop growing once the pH drops to inhibitory levels, usually at or below 5 (48). As a result, commensal oral streptococci co-aggregate with lactate-utilizing bacteria such as *Veillonella* (49). The abundance of not only streptococci but also *Veillonella* sequences in otic secretions is suggestive of similar trophic webs in the middle ear mucosa within microcolonies (5). Such syntrophic partnerships with *Veillonella*, a strict anaerobe, are especially favored within biofilms. Thus, aggregative streptococci are well suited for biofilm growth in the otic mucosa and for syntrophic cooperation with anaerobic partners.

**Fig. 5:**
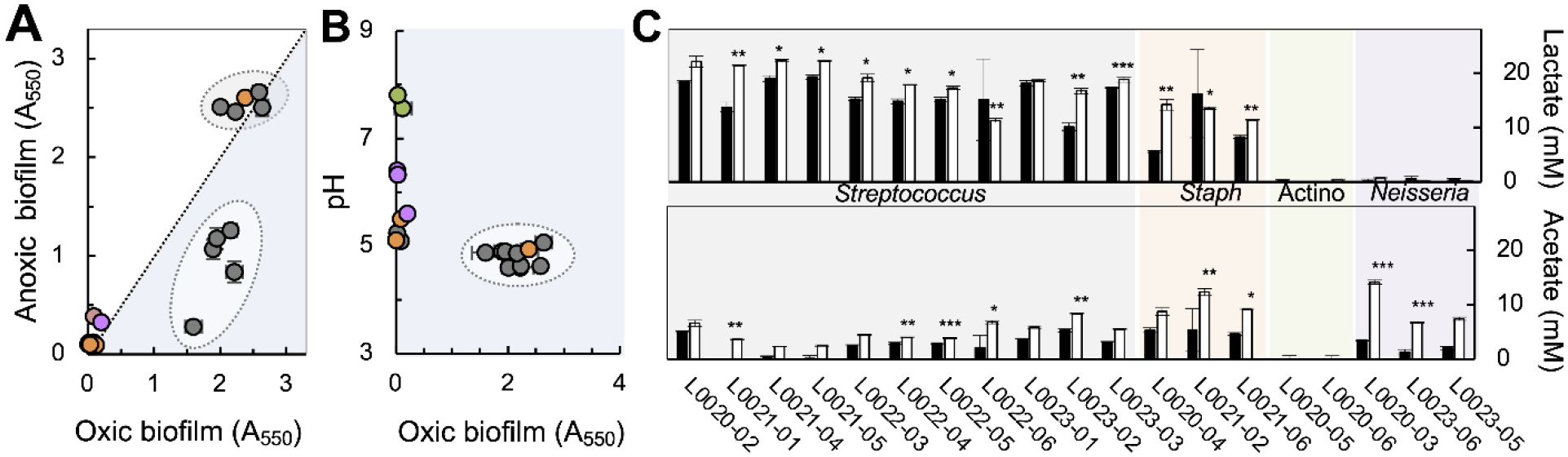
Adaptive responses promoting the establishment of otic trophic webs. (**A**) Biofilm biomass (crystal violet staining, measured as absorbance at 550 nm, A_550_) of otic isolates in oxic (blue) and anoxic (white) cultures. The dashed circles identify two separate clusters of isolates with highest biofilm-forming abilities. (**B**) Correlation between biofilm formation and pH in oxic cultures showing. The circle highlights a cluster of strains with highest biofilm-forming activities and lowest pH. (**C**) Lactate and acetate production (mM) in stationary-phase cultures grown in oxic (black) and anoxic (white) media. The asterisks show significant differences between oxic and anoxic values (*p*≤0.05, *;*p* ≤0.01, **; *p*≤0.001, ***). All data points in **A-C** are average values of three independent biological experiments and are color-coded for *Streptococcus* (gray), *Staphylococcus*(orange), *Neisseria* (purple) and actinobacterial genera *Micrococcus* and *Corynebacterium* (green).

### Antagonistic interactions of otic streptococci with common otopathogens

Commensal oral streptococci mediate intra- and interspecies antagonistic interactions in oral biofilms that are critical to dental and mucosal health (18). Given their oral ancestry, we screened the otic streptococci for their ability to inhibit the growth of known otopathogens (*Streptococcus pneumoniae, Moraxella catharralis*, and non-typeable *Haemophilus influenzae*). For these assays, we followed the same protocol as in other plate assays and spot-plated overnight cultures on TSA plates before incubating them at 37°C for 24 h to allow the colonies to grow. We then covered the plates with a soft (0.7%) agar overlay containing a diluted cell suspension of each otopathogen in a growth medium suitable for their growth. After incubation of the overlayed plates for an additional 24 h, we examined the overlays for zones of growth inhibition on top and around the underlying streptococcal colonies. Growth inhibition in this assay may be the result of nutrient competition, secretion of growth inhibitors or both. Fig. 6 shows representative plate assays for all the otic strains against each otopathogen. Notably, all the otic streptococci inhibited the growth of *S. pneumoniae* and *M. catarrhalis*, producing large zones of clearing beyond the colony edge. The extended region of growth inhibition in the soft-agar overlay is consistent with the secretion of a diffusible inhibitory compound. We also observed antagonistic effects against *H. influenzae*, but they were less pronounced and strain-specific (Fig. 6).

**Fig. 6:**
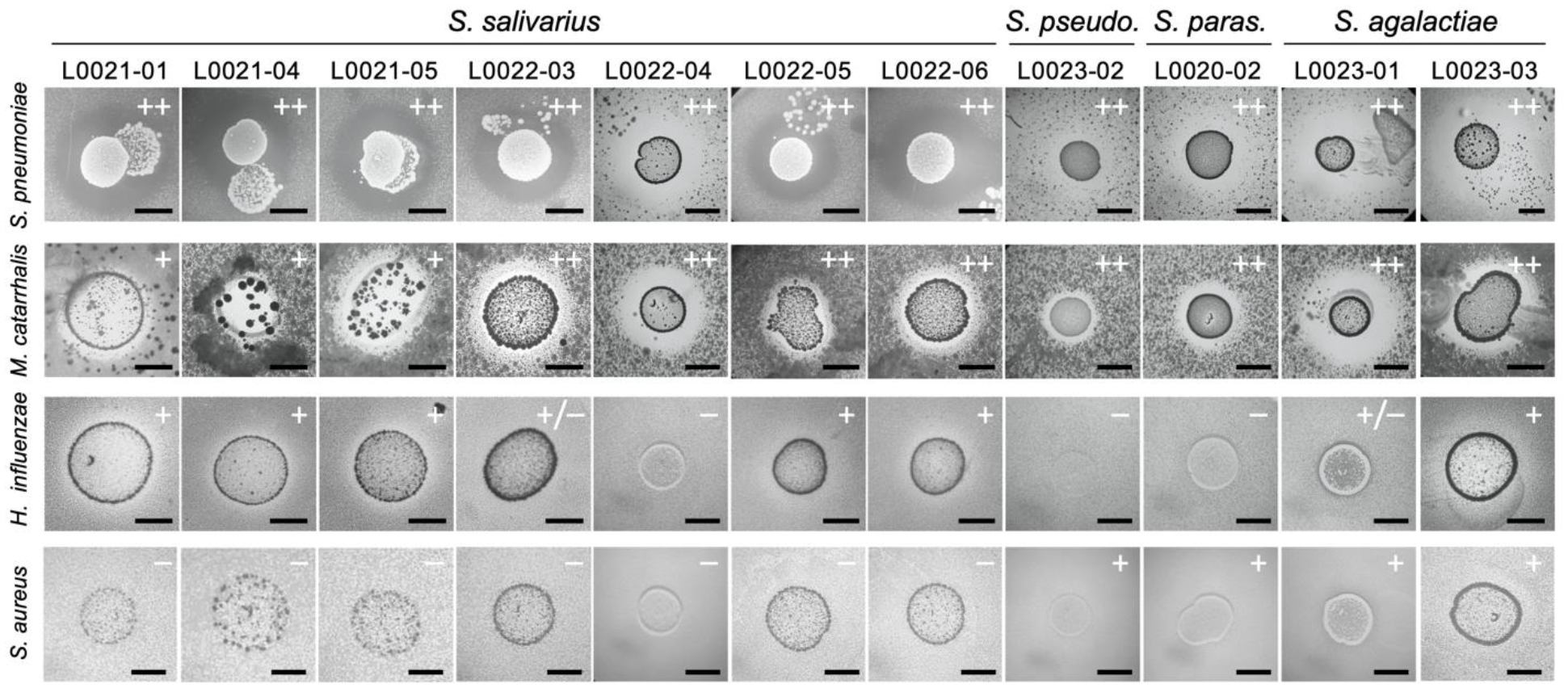
Growth inhibitory effect of otic streptococci against common otopathogens. TSA plates containing 24-h colonies of the otic streptococci were incubated for 24h with soft-agar overlays of the otopathogens *Streptococcus pneumoniae, Moraxella catarrhalis, Haemophilus influenzae* and *Staphylococcus aureus*. The plates show clear areas of growth inhibition of the otopathogen ontop and/or around antagonistic streptococcal colonies underneath (scale bar, 0.5 cm). The symbols indicate average size of the growth inhibition halo around the underlying streptococcal colony in triplicate plate assays (+, <0.4; ++, >0.4; +/-, ∼0.1 but not always reproducible).

We also used the plate assay to screen for potential antagonism of the otic streptococci towards the nasopharyngeal staphylococci that disperse in the aerodigestive tract. As a test strain, we used *S. aureus* subsp. *aureus* JE2 (50), a plasmid-cured derivative of the epidemic community-associated methicillin-resistant *S. aureus* (CA-MRSA) isolate USA300 (51). We observed antagonism by all the non-salivarius isolates (Fig. 6), suggesting species-specific mechanism for growth inhibition by these streptococcal groups (*S. pseudopneumoniae*, *S. parasanguinis* and *S. agalactiae*). The ability of non-salivarius streptococci to inhibit the growth of *S. aureus* is not uncommon. Despite being catalase positive, *S. aureus* is sensitive to hydrogen peroxide produced by *S. pneumoniae* in the nasal mucosa (52). This is because hydrogen peroxide is converted into a highly toxic hydroxyl radical (^•^OH) that rapidly kills *S. aureus* (53). The fact that the closest relatives to the non-salivarius otic streptococci all have catalase-independent mechanisms for anti-oxidative stress resistance (54) and release hydrogen peroxide as a byproduct of their metabolism (55–57) suggests similar mechanisms for interspecies interference with *S. aureus*.

## Discussion

The recovery from otic secretions of close relatives of nasal and oral bacteria (Fig. 2) highlights the role that saliva aerosols play in the dispersal of bacteria throughout the aerodigestive tract. Human saliva carries bacteria shed from oral surfaces such as teeth and gums and spreads them to distant mucosae (58, 59). The constant flux of saliva to the oropharynx (back of the throat) facilitates the formation of aerosols and their carriage to the middle ear every time the Eustachian tube opens and exhaled air is drawn in (5). Not surprisingly, phylogenetic analysis of full-length 16S rRNA sequences recovered from 19 otic cultivars resolved close evolutionary relationships with species that reside or transiently disperse in the oral cavity (Fig. 2). Particularly important were the ancestral ties between the otic streptococci, the most prominent residents of the middle ear communities (5), and pioneer species of oral biofilms (Fig. 2). Most of the otic streptococci were closely related to *S. salivarius,* one of the first colonizers of the human oral cavity after birth and an abundant commensal throughout the life of the host (60). This bacterium disperses in the aerodigestive tract via saliva and forms aggregates that survive stomach passage (61). This allows Salivarius species to enter the small intestine and colonize its mucosa (62). The abundance of *S. salivarius* in saliva also increases its dispersal potential in saliva aerosols, the primary mechanism for seeding of the otic mucosa (5). Aggregation facilitates immunoescape and successful colonization of the otic mucosa. It also promotes coaggregation with anaerobic syntrophic partners and the formation of otic trophic webs (Fig. 1) that mirror those described in oral biofilms. Additionally, oral *S. salivarius* strains mediate antagonistic interactions with virulent streptococci that are critical to prevent tooth decay, periodontal disease, and the spread of respiratory pathogens such as the otopathogen *S. pneumoniae* (63, 64). We see similar interspecies interference of otic *S. salivarius* strains towards common otopathogens (Fig. 6), suggesting similar roles for these middle ear residents in disease prevention.

The non-salivarius otic streptococci were also close relatives of oral species (Fig. 2). For example, one of the isolates (L0020-02) was closely related to *S. parasanguinis*, a bacterium that groups with species of the Mitis group based on 16S rRNA gene sequence analysis and shares with them many of their phenotypic characteristics (27). Like *S. salivarius*, *S. parasanguinis* is one of the early colonizers of the oral cavity (18) and disperses in saliva (65). It produces fimbriae to firmly attach to the biofilms (66), which facilitates its co-dispersal in syntrophic oral aggregates. Also recovered from otic secretions were strains closely related to streptococcal commensals such as *S. pseudopneumoniae* (L0023-02; Viridans group) and *S. agalactiae* (L0023-01 and L0023-03; GBS group). Although the oral ancestors of these otic streptococci have been linked to infective processes in the aerodigestive tract and other body sites (67, 68), the otic relatives readily inhibited the growth of the three most common otopathogens (*S. pneumoniae*, *M. catarrhalis*, and non-typeable *H. influenzae*) (Fig. 6). Furthermore, they were the only otic streptococci that interfered with the growth of *S. aureus* (Fig. 6). Antagonism towards *S. aureus* may involve the production of hydrogen peroxide as a metabolic byproduct, as noted for related oral streptococcal species (55–57). Hydrogen peroxide also functions as a signaling molecule for the co-aggregation of non-salivarius streptococci in syntrophic biofilms (56). Future studies will need to evaluate the role of these streptococcal lineages in producing hydrogen peroxide as a signal for intra and interspecies co-aggregation and as an antagonist of bacterial competitors.

The results presented in this study identified physiological traits of the otic streptococci that could facilitate the colonization of the middle ear mucosa and the formation of syntrophic biofilms. The otic streptococci were temperate swarmers on soft agar plates (Fig. 3), a motile behavior that could allow them to penetrate the otic mucus layer and reach the underlying mucosal epithelium. However, not all the isolates secreted surfactants (Table 2). While endogenous surfactants are often needed for bacteria swarming on semisolid agar surfaces, they are not always needed for efficient swarming through the native mucus layers (20). This is particularly true for bacteria colonizing the middle ear mucosa, where host surfactants abound (69). Furthermore, surfactant production by bacterial colonizers may be undesirable in the otic mucosa as it would change the rheological properties of the mucus and interfere with critical mucosal functions such as antimicrobial activity, immunomodulation and Eustachian tube mechanics (69). The latter is carefully controlled by the secretion of host surfactants that regulate the viscosity and surface tension of the mucus layer (70). This chemical mechanism ensures that the surface tension of the mucus is kept sufficiently low (58 mN/m) to facilitate the rhythmic aperture of the collapsed Eustachian tube and adequate ventilation and decompression of the tympanic cavity (69). Disruption of surfactant homeostasis increases the pressure needed to open the Eustachian tube, risking barotrauma and making the middle ear mucosa more vulnerable to infections (69).

An important finding of our study was the identification of similar phenotypic traits among streptococcal and staphylococcal cultivars recovered from otic secretions that could give them both a competitive advantage during the colonization of the middle ear mucosa. For example, the streptococcal and staphylococcal isolates grew optimally with and without oxygen (Fig. 4A) and secreted mucins and proteases (Table 3), giving them a competitive advantage for growth and reproduction in the middle ear mucosa. Moreover, both groups had fermentative metabolisms and produced lactate (Fig. 5C), a critical intermediate in the syntrophic otic communities (5). This contrasts with otic *Neisseria* (L0020-05 and L0020-06) closely related to *N. flavescens* and *N. subflava*, whose microaerophilic metabolism led to poor growth and extensive flocculation in oxic broth (Fig. 4A). These two *Neisseria* species form a separate clade with members of the family Neisseriaceae that populate the tongue dorsum (71) and although they readily disperse via saliva into the oropharynx (32), they are not positively selected in the middle ear (5). Furthermore, the *Neisseria* cultivars were along with the two actinobacterial isolates (*Micrococcus* spp. L0020-05 and *Corynebacterium* spp. L0020-06) the only cultivars that did not secrete proteases on casein plates (Table 3). Furthermore, they did not form robust biofilms (Fig. 5A) nor did they produce fermentative byproducts such as lactate (Fig. 5C), a key metabolic intermediate in the otic trophic webs (Fig. 1). Not surprisingly, despite being the most abundant nasal phylum (38), actinobacterial sequences are poorly represented in otic secretions (5).

The most notable difference between the staphylococcal and streptococcal isolates was, however, the co-aggregative behavior of most *Streptococcus*, which is critical for the formation of microcolonies with anaerobic syntrophic partners (Fig. 1). Aggregative behavior may in fact be the most successful strategy of oral streptococci during the colonization of the middle ear mucosa. Most of the streptococcal isolates autoaggregated when growing on agar-solidified media (Fig. 3B), a phenotype associated with increased resistance to antimicrobials and immunoescape (72). Aggregation allows oral streptococci to recognize and partner with other bacteria during the formation of biofilms (49). Thus, streptococci coaggregate with actinomyces to colonize the tooth surface and recruit other bacteria during the formation of the dental plaque (18, 73). Metabolic interactions between lactic acid-producing strains of *Streptococcus* and *Veillonella* spp., which ferment lactate, are critical for coaggregation during the early stages of biofilm formation on oral surfaces (49). Fusobacteria mediate early coaggregation as well, forming physical bridges across the biofilms and promoting the attachment of non-coaggregating aerobes and anaerobes (18, 73). These syntrophic interactions sustain the growth of the dental plaque throughout all dentition stages and the formation of subgingival biofilms in the predendate and postdentate states (59). The widespread presence and abundance of syntrophic co-aggregates in the oral cavity promotes their co-dispersal in saliva (74) and provides a mechanism for their co-immigration in saliva aerosols. These coaggregates readily establish otic trophic webs (Fig. 1) similar to those described in oral biofilms (5). Consistent with this, all the streptococcal isolates characterized in this study were highly aggregative (Fig. 4), formed robust biofilms (Fig. 5A) and produced lactate as primary byproduct of their fermentative metabolism (Fig. 4C). These adaptive traits allow streptococci to grow and reproduce in the middle ear mucosa with obligate anaerobic, syntrophic partners such as *Prevotella, Fusobacterium* and *Veillonella* (5). The syntrophic microcolonies metabolize and ferment host mucins and proteins in the otic mucosa (Fig. 1), indirectly controlling the viscoelastic properties of the mucus layer and Eustachian tube functionality (69). The detection of a differential gradient of mucin gene expression along the tympanic cavity and Eustachian tube (69) suggests a high degree of spatial heterogeneity in bacterial colonization as well. Shaped like an inverted flask (4), the posterior region of the Eustachian tube is more readily seeded with saliva aerosols during the cycles of tubal aperture. Concentration of streptococcal aggregates in this region closer to the nasopharyngeal opening of the Eustachian tube could provide increased protection against otopathogens, which typically reside in nasal reservoirs. Future research should therefore consider the mechanisms that allow otic streptococci to co-aggregate with syntrophic partners, their spatial distribution in the otic mucosa and antagonistic interactions with transient migrants. This knowledge is important to understand the functionality of the otic communities and how they influencehost functions and the outcome of infections.

## Methods

### Bacterial strains and culture conditions

The bacterial strains used in this study include 19 cultivars isolated from otic secretions (5). Briefly, the samples were collected with a single swab from the left and right nasopharyngeal openings of the Eustachian tube in 4 young (19-32 years old), healthy adults recruited as part of a larger study approved by the Michigan State University Biomedical and Physical Health Review Board (IRB # 17-502). The cultivars were isolated as single colonies on Tryptic Soy Agar (TSA) plates (30g/L of Tryptic Soy Broth from Sigma Aldrich and 15g/L of Bacto Agar from BD) grown at 37°C. The isolates were routinely grown overnight in 5 ml of Tryptic Soy Broth (TSB) at 37°C with gentle agitation. For growth studies, we transferred mid-log phase (OD_600_ ∼0.5) TSB cultures twice (initial OD_600_ of 0.1) to prepare a stationary phase (∼0.9-1.0 OD_600_) inoculum for growth assays in Corning® 96- well clear round bottom TC-treated microplate (Corning 3799). Growth was initiated with the addition of 18 µl of the inoculum to 162 µl of TSB per well and monitored spectrophotometrically every 30 min (OD_630_ readings after 0.1 sec of gentle agitation) while incubating the plates at 37°C inside a PowerWave HT (BioTek) plate reader. Each plate contained a well with TSB medium without cells to use as a blank. Growth in anoxic medium was monitored in a similar way but in a plate reader housed inside an 855-ABC Portable Anaerobic Chamber (Plas Labs, Inc.) containing a headspace of N_2_:CO_2_ (80:20).

### DNA sequencing and phylogenetic analyses

For taxonomic and phylogenetic analyses, we grew 19 otic isolates (Table 1) in 2 ml of TSB at 37°C for 24 h and harvested the cells by centrifugation (25,000 x *g* for 5 min) in an Eppendorf 5417R refrigerated centrifuge prior to extracting the genomic DNA with a FastDNA^TM^ Spin kit (MP Biomedicals). Library preparation with an Illumina Nextera kit and whole genome sequencing in an Illumina NextSeq 550 platform were at the Microbial Genome Sequencing Center (MiGS; Pittsburgh, PA). We used the FastQC tool from the Babraham Institute (https://www.bioinformatics.babraham.ac.uk/projects/fastqc/) for sequence quality control and Trimmomatics (75) for cleaning/trimming of the Illumina short reads. After assembling the genomes *de novo* with the Spades assembler (76), we identified the 16S rRNA gene sequences in the contigs with the BAsic Rapid Ribosomal RNA Predictor (Barnap) (https://github.com/tseemann/barrnap). The 16S rRNA gene sequences were deposited in the GenBank database under individual accession numbers (Table 1). We used the full-length 16S rRNA sequences to identify the closest species (% identity) in the GenBank database using the nucleotide Basic Local Alignment Search Tool (BLAST) at the U.S. National Center of Biological Information (NCBI) using an identity species cutoff value of 98.7% (23). We retrieved the 16S rRNA gene sequences from the closest type strains listed in the SILVA rRNA database (https://www.arb-silva.de) and aligned them to the otic sequences with the MUSCLE program in the MEGA X software (77). We used the alignment to build a maximum-likelihood phylogenetic tree and calculate bootstrap confidence values for each node using 1,000 replications. The tree shows bootstrap values above 50% (78).

### Catalase assay

Frozen stocks of the otic isolates were directly streaked on 1.5% (w/v) TSA plates to grow individual colonies at 37°C overnight. We spread each colony onto a microscope slide and added a drop of freshly prepared 3% hydrogen peroxide. Catalase-positive strains breakdown the hydrogen peroxide into water and oxygen gas, which generates bubbles. Lack or weak production of bubbles is used to designate a strain as catalase-negative.

### Swarming motility and surfactant detection assays

We screened each otic isolate for their ability to move on soft (0.5% and, when indicated, 0.4% w/v agar) TSA plates, as a modification of a previously described assay (79). For these assays, we first grew each isolate and the positive control (*P. aeruginosa* PA01) in TSB at 37°C overnight (OD_600_ ∼1) and prepared a diluted TSB inoculum (OD_600_ 0.1). We pipetted a 5-µl drop of the diluted culture onto the surface of the soft agar plates and allowed it to absorb until completely dry (∼30 min). We then incubated the plates at 37°C and photographed the areas of growth at 18, 42 and 62 h against a ruler using a dissecting scope (Leica MZ6) at a magnification of 0.8X and 1X. The photographs were then analyzed with the ImageJ software (80) to measure the colony diameter over time and calculate the area expansion (swarming distance) from the initial inoculation spot.

We also screened the ability of the otic isolates to produce surfactants with a previously described atomized oil assay (40). For this, we plated a 5-µl drop of the diluted TSB culture (OD_600_ of 0.1) on agar-solidified (1.5% w/v) TSA medium, allowed the inoculum to absorb for ∼30 min, and incubated the plates at 37°C for 24 h. Using an airbrush (type H; Paasche Airbrush Co., Chicago, IL), we applied a fine mist of mineral oil onto the plate surface. Surfactant-producing colonies readily display a halo of mineral oil dispersal whose size provides a semiquantitative measure of surfactant secretion (40). Photography and halo diameter visualization was as described above for swarming assays, except that we measured the size of the oil dispersal zone from the colony edge. All strains were tested in three independent swarming and surfactant assays plates to calculate the average and standard deviation values.

### Protease and mucinase plate assays

We used TSA plates containing 5% lactose-free, skim milk (Fairlife, LLC) or 0.5% Type II porcine gastric mucin (Sigma Aldrich) to screen the otic isolates for mucinase and protease secretion, respectively, using *P. aeruginosa* PA01 as a positive control. For these assays, we spot-plated 5 µl of overnight TSB cultures and incubated at 37°C for 24 h, as described earlier for the surfactant assays. Strains that secrete proteases to the medium degrade the milk’s casein and produce a clear halo around the area of growth after 24 h of incubation. Mucinase producers have zones of mucin lysis around or under the colony that show as zones of discoloration after staining with 7ml of 0.1% amido black for 30 min and destaining with 14 ml of 2.5 M acetic acid for 30 min. When indicated, plates were incubated for 48 h to confirm emerging phenotypes. Each strain was tested in triplicate plates and each was photographed on a lightboard (A4 LED Light Box 9×12 Inch Light Pad, ME456) with an iPhone 11 at 2.4x magnification.

### Organic acid detection in culture supernatant fluids

We grew triplicate stationary phase cultures of the otic isolates in oxic and anoxic TSB medium at 37°C and harvested the culture supernatant fluids by centrifugation (14,000 rpm, 10 min). We measured the pH of the supernatant fluids (5 ml) with a pH probe (Thermo Scientific^TM^ Orion^TM^ 720A+ benchtop pH meter) and stored 1 ml of the samples at –20°C for chemical analyses by high performance liquid chromatography (HPLC). Once thawed, we filter-sterilized 250 µl of the supernatant fluid into 1-ml HPLC vials and measured their organic acid content in a Shimadzu 20A HPLC equipped with an Aminex HPX-87H column and a Micro-Guard cation H^+^ guard column (Bio-Rad, Hercules, CA) at 55°C, as previously described (81). As controls, we included samples with TSB medium and standard solutions of acetate, lactate and pyruvate (provided at 1, 2, 5, 10 and 20 mM concentrations).

### Biofilm assays

We used a previously described assay (82) to test the ability of the otic cultivars to form biofilms in Corning® 96-well clear round bottom TC-treated microplates (Corning 3799). We first grew overnight cultures in TSB with gentle agitation (∼200 rpm) and used them to prepare a diluted cell suspension (OD_600_ ∼0.1) for inoculation (18 µl) into TSB medium (162 µl per well). Each isolate was tested in 8 replicate wells. After incubating the plates at 37°C for 24 h, we removed the planktonic culture, washed the wells with ddH_2_O and stained the surface-attached cells with 0.1% (w/v) crystal violet. We then rinsed the wells with water and let the stained biofilms to dry overnight at room temperature before solubilizing the biofilm-associated crystal violet with 180 µl of 30% glacial acetic acid and measured the crystal violet in the solution spectrophotometrically at 550 nm (82).

### Growth inhibition plate assays

We screened the otic streptococcal isolates for their ability to inhibit the growth of bacterial species (*S. pneumoniae, M. catarrhalis,* and non-typeable *H influenzae*) commonly associated with infections of the middle ear (83). As test strains, we used *S. pneumoniae* ATCC 6303 and *M. catarrhalis* ATCC 25238 (from the laboratory strain collection of Dr. Martha Mulks, Department of Microbiology and Molecular Genetics, Michigan State University) and a non-typeable *H. influenzae* (NTHi) strain isolated by Dr. Poorna Viswanathan in the teaching lab of the Department of Microbiology and Molecular Genetics (Michigan State University). The NTHi strain was confirmed prior to experimental use by multiplex PCR confirmation, as described previously (84). We also included for testing the laboratory strain *S. aureus* JE2, which was kindly provided by Dr. Neal Hammer (Department of Microbiology and Molecular Genetics, Michigan State University). The otic streptococci and *S. aureus* JE2 were routinely grown in 5 ml TSB at 37°C with gentle agitation to prepare overnight cultures for the plate assays. *S. pneumoniae* and *M. catarrhalis* were grown at 37°C overnight in 5 mL of brain heart infusion (BHI) broth (Sigma-Aldrich) without agitation. The NTHi reference strain of *H. influenzae* was also grown statically at 37°C but in supplemented BHI (sBHI) (85), which contains (per L): 30 g BHI, 0.01 mg hemin (Bovine, Sigma Aldrich), and 0.002 mg β- Nicotinamide adenine dinucleotide sodium salt (Sigma Aldrich). All incubations were in a 37°C incubator with a 5% CO_2_ atmosphere except for *S. aureus*, which were in air.

We used the spot-on-lawn method (86) to investigate antagonistic interactions between the otic streptococci and test strains. We first spotted 5 µl of a diluted (OD_600_ 0.1) overnight culture of each streptococcal strain onto a 1.5% (w/v agar) TSA plate and allowed it to dry for 30 min at room temperature before incubating at 37°C for 24 h to grow the colonies. We then overlayed the plates with a warm (55°C) 8-ml layer of soft-agar (0.7%, w/v, final concentration) medium (TSA, BHI or sBHI) containing the test strain (OD_600_ 0.1). The general procedure to make 0.7% agar overlays was to autoclave 6 ml of 1% agar-solidified growth medium, cool down the melted agar in a 55°C water bath, add 2 ml of the test strain culture to a final OD_600_ of 0.1, and mix by inversion before pouring over the TSA plate surface with the otic colonies. To make sBHI overlays, we added the chemical supplements to 6 ml of warm (55°C), melted 1% (w/v) agar BHI before mixing with 2 ml of an overnight NTHi culture to a final OD_600_ of 0.1. The overlays were allowed to solidify at room temperature before incubating for an additional 24 h at 37°C in an incubator with or without (*S. aureus* overlay) 5% CO_2_. These culture conditions promoted the growth of the test strains as a turbid lawn in the overlays after 24 h, except for areas of growth inhibition (halos or clear zones) on top and around colonies of antagonistic streptococci growing underneath. At the end of the incubation period, we photographed the overlayed plates with a dissecting scope (0.63x objective) against a ruler and used the ImageJ program (4) to measure the size of the growth inhibition zone from the streptococcal colony edge underneath in triplicate biological replicates.

## Acknowledgements

The authors would like to thank Dr. Michaela TerAverst and Nicholas Tefft at Michigan State University for assistance with the HPLC analyses and Drs. Batsirai Mabvakure and Heather Blankenship at the Michigan Department of Health and Human Services Bureau of Laboratory for guidance on Illumina short read assembly and contig gene analysis. We are also thankful to Drs. Martha Mulks, Poorna Viswanathan and Neal Hammer for providing the test strains used in the growth inhibition plate assays.

